# Data-driven multiscale modeling reveals the role of metabolic coupling for the spatio-temporal growth dynamics of yeast colonies

**DOI:** 10.1101/344226

**Authors:** Jukka Intosalmi, Adrian C. Scott, Michelle Hays, Nicholas Flann, Olli Yli-Harja, Harri Lähdesmäki, Aimée M. Dudley, Alexander Skupin

## Abstract

**Motivation:** Multicellular entities, such as mammalian tissues or microbial biofilms, typically exhibit complex spatial arrangements that are adapted to their specific functions or environments. These structures result from intercellular signaling as well as from the interaction with the environment that allow cells of the same genotype to differentiate into well-organized communities of diversified cells. Despite its importance, our understanding on how cell–cell and metabolic coupling produce functionally optimized structures is still limited.

**Results:** Here, we present a data-driven spatial framework to computationally investigate the development of one multicellular structure, yeast colonies. Using experimental growth data from homogeneous liquid media conditions, we develop and parameterize a dynamic cell state and growth model. We then use the resulting model in a coarse-grained spatial model, which we calibrate using experimental time-course data of colony growth. Throughout the model development process, we use state-of-the-art statistical techniques to handle the uncertainty of model structure and parameterization. Further, we validate the model predictions against independent experimental data and illustrate how metabolic coupling plays a central role in colony formation.

**Availability:** Experimental data and a computational implementation to reproduce the results are available at http://research.cs.aalto.fi/csb/software/multiscale/code.zip.

## 1 Introduction

Multicellular organisms and colonies of unicellular microbes are able to form characteristic structures. While it is generally accepted that the structure and functions of tissue and organs are genetically encoded, more recently it has been demonstrated that the morphologies of biofilms like *Saccharomyces cerevisiae* yeast colonies have a strong genetic component (Stovivcek *et al.*, 2010; Granek *et al.*, 2013; Taylor and Ehrenreich, 2014). Together with the frequently observed growth medium dependency of yeast cultures this underpins the importance of genome-environment interactions for phenotype development (Zahn and Purnell, 2016). To understand the underlying mechanisms, it is key to investigate how metabolic coupling is influencing individual cell states and instructing structure formation.

Yeast colonies represent an efficient experimental model system to investigate how metabolic dynamics and spatial coupling determine morphogenesis because yeast exhibits cell state transition in dependence on the environment, fast growth, and can be easily genetically modified (Fig. 1A). Although yeast is a unicellular organism, it can form rather complex colony structures in a strain specific manner (Vachova *et al.*, 2009, 2011). Recently, we have shown that predominant changes in morphology from smooth to wrinkled “fluffy” structures can be induced by aneuploidy as a multicellular phenotype switch (Tan *et al.*, 2013). Despite the systematic genetical characterization of this switch, the question how the gain or loss of a chromosome copy leads to a significant change in morphology is not understood.

**Figure 1:**
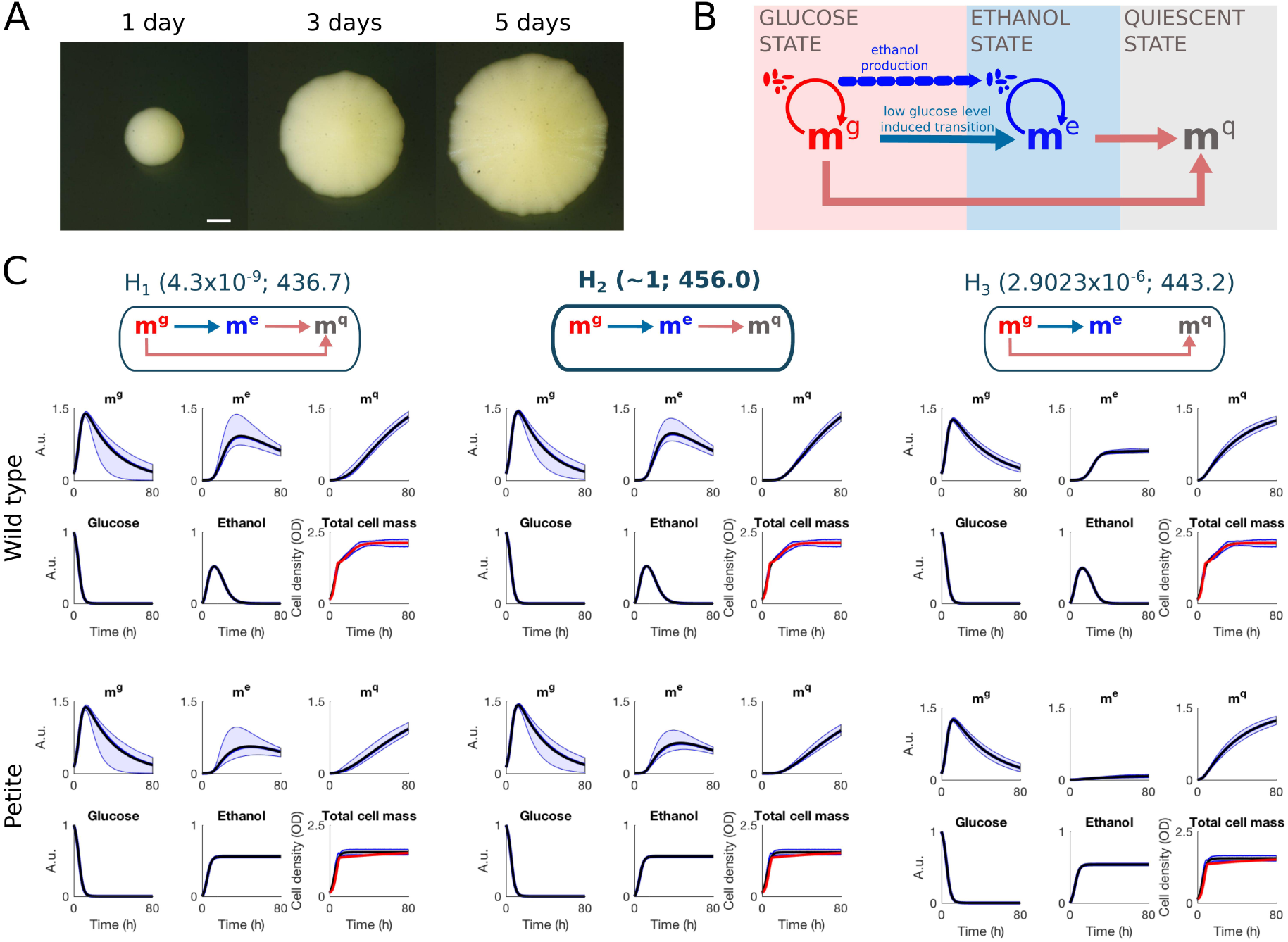
Illustration of real colony growth and summary of microenvironment model inference. (A) A real colony growing on a nutrient rich agar. (B) Schematic illustration of the microenvironment model. (C) Illustration of the alternative metabolic switching routes (hypotheses *H*_1_, *H*_2_, and *H*_3_) and summary of microenvironment model inference. The hypothesis *H*_1_ contains both possible transitions from the glucose state to the quiescent state and the hypotheses *H*_2_ and *H*_3_ can be obtained by removing one of the routes (these hypotheses correspond to setting the switching rate parameters *β*_2_ and *β*_3_ in the model to zero, respectively). Each hypothesis is accompanied with the posterior probability and the estimated logarithmic marginal likelihood (shown in parentheses after hypothesis). The estimated marginal posterior predictive distributions are illustrated using 99% quantiles (light blue region) as well as mean (black line) and median (blue line). The experimental data (total cell mass) is illustrated using red color.

Mathematical modeling can provide essential insights into the underlying processes as it allows quantitative investigation of the coupling between metabolic and spatial growth dynamics. A general challenge is thereby to cover and parameterize the relevant scales ranging from intra- and intercellular interactions to population and environment dynamics. Existing multiscale modeling approaches for complex multicellular systems typically rely on large sets of physiological parameters that are often not easily accessible in experiments (Kang *et al.*, 2014; Doloman *et al.*, 2017). Other spatiotemporal modelling approaches are based on homogeneity assumption and simulate partial differential equations neglecting the discrete properties of cells. While being useful in building a general understanding of different mechanisms across the scales, most of these approaches do not allow experimentally-based model construction and validation. Experimentally-based model construction approaches have been successfully applied in the context of mechanistic modeling of molecular mechanisms (Schulz *et al.*, 2009; Intosalmi *et al.*, 2015; Chan *et al.*, 2016) and extending these approaches to more complex multiscale models will be essential for methodological advancement in systems biology (Skupin *et al.*, 2010).

Here, we develop a new multiscale modeling framework for yeast colony formation. In contrast to previous approaches that simulate individual cells (Walther *et al.*, 2004), our framework is based on an approximation that discretizes the spatial domain into elementary cubes and allows us to model the microenvironment dynamics under the homogeneity assumption. Further, the elementary cube approximation enables us to model the information flows (like nutrient transport or the flow of signaling molecules) and mass transfer (movement of the growing cell mass) by means of computationally efficient flux mechanisms.

To construct a growth and cell state model for homogeneous microenvironment dynamics, we combine ordinary differential equation (ODE) modeling with experimental data using advanced statistical techniques and, by means of this objective approach, learn the metabolic switching mechanisms as well as the corresponding model parameterization directly from the data. The calibrated microenvironment model is then embedded into the spatial framework and, once the spatial model is calibrated using colony morphology data, we are capable of predicting the cell mass, cell state, nutrient, and metabolic distributions throughout the colony formation process.

Our model construction process utilizes measurements from two different yeast strains. First, we calibrate the model using time-course data from wild-type yeast cells (YAD145) and subsequently the calibrated model is validated against independent measurements from a respiratory deficient (petite) yeast strain (YAD479). These genotypically different training and validation strains are known to result in distinct colony morphologies and therefore the validation approves that our multiscale model captures essential mechanisms across the scales spanning from microenvironment dynamics to the spatiotemporal colony formation dynamics.

## 2 Methods

### 2.1 Growth curve data

The experimental procedures are detailed elsewhere (Tan *et al.*, 2013). In brief, growth curves in suspension of the FY4 (YAD145) strain and its petite version YAD479 (that is unable to metabolise nonfermentable carbon sources like ethanol) were measured on a TECAN Sunrise (Tecan). Initially, 200 µL were seeded from running cultures at 5 × 10^5^ cells/mL and cell number was monitored by optical density (OD) every 15 min for 88 hours in YPD medium containing 2% glucose at 30°C. The growth data are provided in a machine readable format as a part of the computational implementation and details about preprocessing can be found in Supporting Information (Fig. S1).

### 2.2 Colony footprint area data

Colony formation was measured by our custom built colony imaging system (Scott and Dudley, unpublished results). Colonies from single cells were grown on YPD-agar plates with 5% glucose for 7.5 days in an incubator at 30°C, and photographed every 20 minutes. Colony areas were extracted from each image by a script for NIH ImageJ (Schneider *et al.*, 2012) (See Supplementary Information for details on image capture and analysis).

### 2.3 Bayesian techniques for ODE model calibration

The parameters and structure of ODE models are calibrated within the Bayesian framework (see e.g. Girolami, 2008). In brief, we link the model output with time-course data *D* via the likelihood function 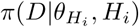 where 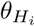 is the parameter vector under the hypothesis *H_i_* about the model structure (*i* = 1,…, *n*). A Bayesian statistical model can be constructed by combining the likelihood function with a prior distribution over the parameters, 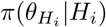 (Robert, 2007). Bayes’ theorem yields the parameter posterior distribution 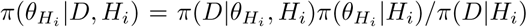, where *π*(*D*|*H_i_*) is the marginal likelihood. The marginal likelihoods *π*(*D*|*H_i_*) can be used to compute the posterior distribution over the hypotheses, i.e. 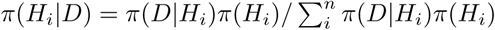, where *π*(*M_i_*) is the prior distribution over the alternative models. In this study, the prior distribution over the alternative hypotheses is assumed to be uniform.

### 2.4 Population-based Markov chain Monte Carlo sampling

Neither the posterior distributions nor the marginal likelihoods can be analytically solved for our models and, consequently, the posterior analysis needs to be carried out using numerical techniques. For this purpose, we use the population-based Markov chain Monte Carlo (MCMC) sampling and thermodynamic integration (Jasra *et al.*, 2007; Calderhead and Girolami, 2009).

To implement a population-based Markov chain Monte Carlo sampler, we consider a product form of the target density

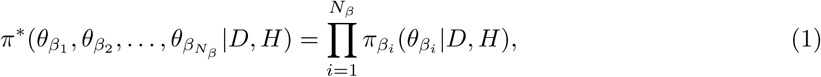

where 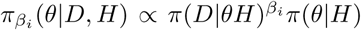 is the power posterior for fixed temperatures 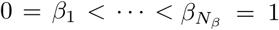 (Jasra *et al.*, 2007; Calderhead and Girolami, 2009). The distributions 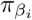, including the posterior distribution *π*(*D*|*θ, H*)*π*(*θ*|*H*), are marginal distributions of the product form of the target density. By means of population-based MCMC sampling, we draw samples from the individual marginal distributions as well as allow global moves between neighboring temperatures (for details, see Jasra *et al.* (2007); Calderhead and Girolami (2009)).

In this study, we select the temperatures according to the formula

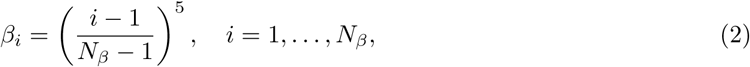

and use altogether 30 temperatures (*N_β_* = 30). Before running the sampler, we use local gradient-based deterministic multistart optimization to determine the highest peak in each temperature and the corresponding points are then used as an initial state for the sampler. For the multistart optimization, we use our own optimization routine which is implemented in Matlab according to the guidelines given in references Raue *et al.* (2013, 2015). The actual sampling is run in two parts. First, 10^5^ samples are drawn so that the normal proposal distributions are adaptively tuned based on the estimated covariance of the previous 7500 samples. After this burn-in and adaption period, the proposal distributions are fixed and every 1000th sample is collected until 2500 samples are obtained. We run four independent samplers under each alternative hypothesis and the convergence of the chains is monitored by means of the potential scale reduction factors (Gelman *et al.*, 2013) and visual inspection over all temperatures. After checking the convergence, the samples from four independent runs are combined and the posterior analysis is carried out using all 10^4^ samples.

### 2.5 Bayesian optimization

The parameters of the spatial model are optimized by using the Bayesian optimization technique which is tailored for global optimization of cost functions (Jones *et al.*, 1998; Ghahramani, 2015).

To calibrate the spatial model, we need to minimize a target function 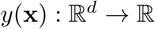 with respect to the parameters **x** (we note here that this notation applies only to this subsection). The evaluation of the target function is computationally costly and, to be able to find the minimum using as few as possible function evaluations, we approximate *y*(**x**) by means of a Gaussian process *f*(**x**). Formally, we can write

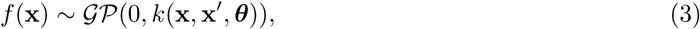

where

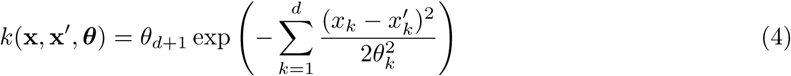

is the squared exponential kernel function and 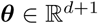 is a parameter vector (for details about Gaussian processes, see e.g. Rasmussen and Williams (2006)). We assume that the approximation error is normally distributed i.e.

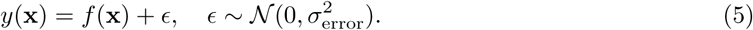

Based on the above definitions, the prior distribution for the approximated function values *f_n_* = *f*(**x***_n_*), *n* = 1,…, *N* is the zero-mean multivariate normal distribution, i.e.

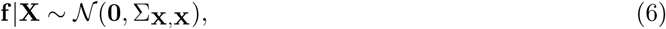

where **f** = [*f*(**x**_1_), *f*(**x**_2_),…, *f*(**x***_N_*)]′, **X** = [**x**_1_, **x**_2_,…, **x***_n_*], and {Σ**_X_**,**_X_**}*_ij_* = *k*(**x***_i_,* **x***_j_, **θ***)*, i, j* = 1,…, *N*. It follows also that

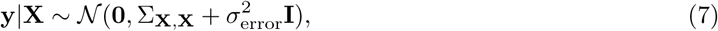

where we have used the above notation, **y** = [*y*(**x**_1_)*, y*(**x**_2_),…, *y*(**x***_N_*)]′, and **I** is the identity matrix. The marginal likelihood is 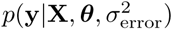 where we have explicitly added the kernel parameters ***θ*** and error variance 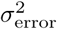 to emphasize that the distribution and the marginal likelihood depend on this parameterization.

Given a set of evaluated function values at certain points (i.e. **y** = [*y*(**x**_1_)*, y*(**x**_2_),…, *y*(**x***_N_*)]′), we can generate a probabilistic prediction on the function value *y*(**x**\*) at an arbitrary point **x**\* in the domain. The prediction about the function value *y*(**x**\*) can be generated in a form of a random variable *y*\* which follows the joint distribution in Eq. 7. By conditioning *y*\* on the evaluated values, we obtain

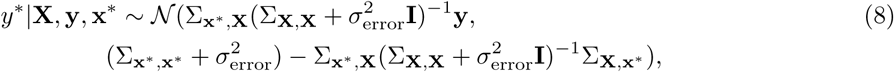

where 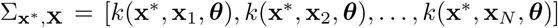, 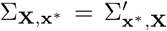, and 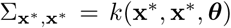. The probabilistic nature of the prediction makes it also possible to predict the next point at which it is most beneficial to evaluate the function value in the context of minimization problem (Jones *et al.*, 1998). The optimal evaluation point can be chosen by finding the point **x**\* which maximizes the expected improvement function

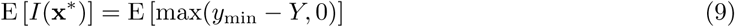

where *y*_min_ is the minimum of the evaluated function values so far and *Y* = *y*\*|**X**, **y**, **x**\* (for details and illustrative examples, see e.g. Jones *et al.* (1998)). The expected improvement (Eq. 9) can be expressed in the closed form

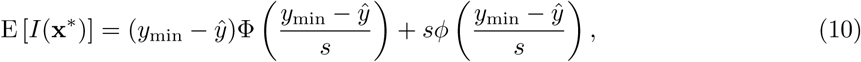

where *ϕ* and Φ are the standard normal density and distribution function, respectively, and 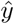 and *s* are the mean and standard deviation of the normal distribution in Eq. 8, respectively (Jones *et al.*, 1998).

The actual optimization routine consists of two steps: (1) fitting the response surface by maximizing *p*(**y**|**X**) (Eq. 7) with respect to the hyperparameters 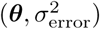 and (2) finding the optimal point for next function evaluation by maximizing the expected improvement (Eq. 9). The steps are carried out sequentially and the response surface is always fitted using a set of evaluated function values which are standardized to have a zero mean and standard deviation of one. In our implementation, the hyperparameters of the Gaussian process model and the next evaluation point with respect to the expected improvement are optimized using fminunc and fmincon optimization routines in Matlab, respectively. The hyperparameter optimization is initialized using parameter values *θ*_1_ = *θ*_2_ = *θ*_3_ = 1*, σ*_error_ = 0.1 which correspond to a smooth Gaussian process response surface. In the context of expected improvement optimization, we utilize a multistart optimization strategy for which the initial points are obtained by means of Latin hypercube sampling (lhsdesign function in Matlab). The sequential procedure is repeated until the expected improvement goes under a threshold (10^−46^ in this study) or the maximum number of iterations of steps (1) and (2) is reached.

### 2.6 Formal definition of the spatial framework

We discretize the space by dividing it into finite size elementary cubes each having a constant volume (for illustration, see Fig. 2). The cubes are indexed by their location in a 3D array i.e. mass in different metabolic states at different spatial locations can be expressed by writing

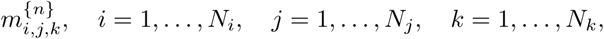

where {*n*} ∊ {g,e,q} denotes the metabolic state. The total mass at each location can be computed by summing the cell masses in distinct metabolic states, i.e.

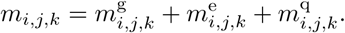

The cubes interact through their fill levels and the cell mass is flowing from a high concentration to a low concentration once a certain threshold is exceeded. The amount of mass exceeding the threshold can be interpreted as pressure that pushes the cell mass onwards. This pressure is computed based on a thresholded total mass distribution over the space. The thresholded total mass at a certain spatial location is defined to be

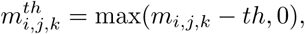

where *th* is the threshold parameter.

#### 2.6.1 Mass movement

In context of mass movement modeling, it needs to be taken into account that the moving cell mass carries along fractions of cells in different metabolic states. The fractions carried along can be taken to be proportional to the cell state fractions in the source cubes (the cubes from which the mass is moved). Consequently, it is natural to model the mass movement by means of the equation

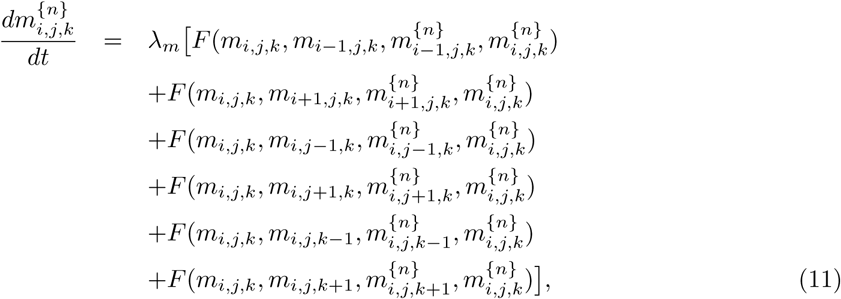

where *λ_m_* is the mass movement rate parameter,

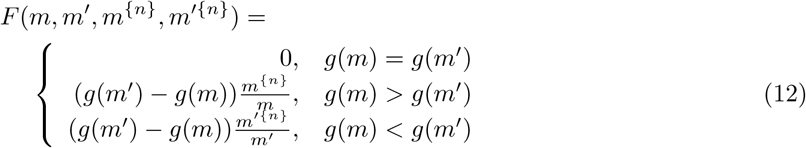

and *g*(*m*) = max(*m* − *th,* 0) is a function which takes care of the thresholding with the parameter *th*. At the agar-cell mass interface, the mass movement into the agar is prevented by mapping the corresponding values of the function *F* to zero.

To show that the mass is conserved through the movement, we can consider mass movement between two elementary cubes *m* to *m*′. Based on our model structure, we have

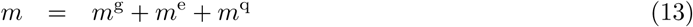

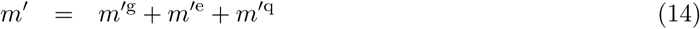

and the thresholded total cell masses in these two cubes are

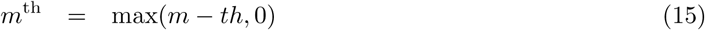

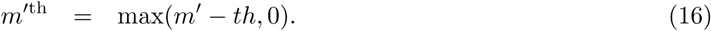

Without losing any generality, we can assume *m*^th^ > *m*′^th^. Now

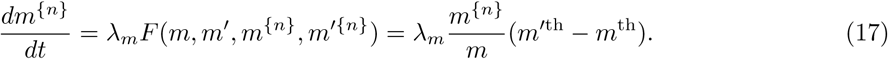

and

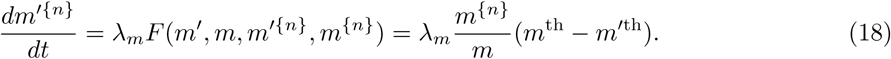

From Eqs. 17 and 18, we can deduce

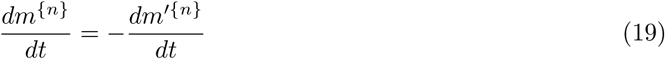

which shows that the mass is conserved through the movement. Because the net mass movement defined in Eq. 11 is a sum of six pairwise movements, the mass is conserved also through the net movement.

#### 2.6.2 Nutrient transfer

The nutrient transfer can be described in a similar manner as the mass movement but, in this context, we do not need to threshold the distribution because the nutrient diffusion can be seen to occur freely in the media. Further, nutrient transfer can be simply defined using fluxes between the neighboring cubes (in the context of mass movement, we needed to take the fractions of different cell types into account). If we consider the nutrient concentrations *n_i,j,k_*, *i* = 1,…, *N_i_*, *j* = 1,…, *N_j_*, *k* = 1,…, *N_k_*, the nutrient transfer can be described using the equation

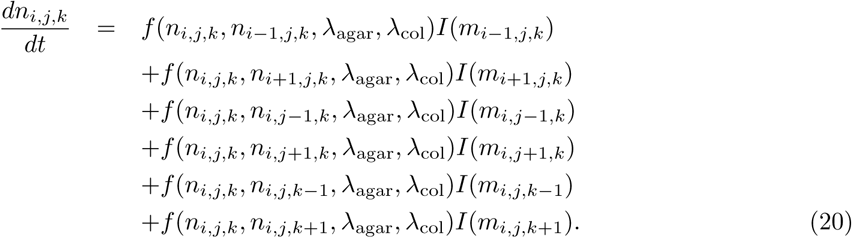

Here,

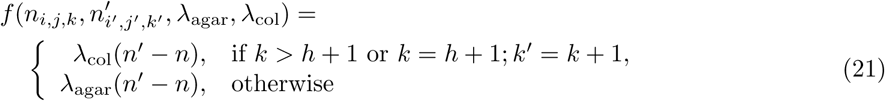

where *λ*_col_ and *λ*_agar_ are the nutrient transfer rate parameters within the colony and agar, respectively, and *h* is the height of the agar given as the number of elementary cube layers. Further, the domain in which the nutrient transfer takes place is determined by the indicator function

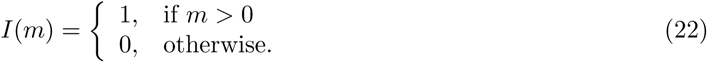

In other words, the mass distribution dependent domain for the nutrient transfer consists of the cubes which have a positive cell mass concentration.

### 2.7 Computational implementation

Mathematical models, population-based MCMC sampler, and Bayesian optimization were implemented in Matlab (The MathWorks Inc., Natick, MA, USA). ODE systems were solved using the ode15s solver and the full multiscale model was simulated using the Euler method with a time-step of 0.0025 h.

## 3 Results

### 3.1 Dynamic model for cell growth and metabolic switching in homogeneous medium

Depending on external conditions and intracellular state, yeast cells can either metabolize glucose or ethanol for growth or remain in the so-called quiescent state. The diauxic shift between the different metabolic states is determined by nutrient sensing pathways and if the extracallular glucose level becomes low, cells change their metabolic wiring towards a state that allows growth on ethanol produced during growth on glucose (DeRisi *et al.*, 1997; Galdieri *et al.*, 2010). Cells can also switch to a quiescent state in which they act as passive by-standers that do not grow nor produce any aromatic alcohols. The metabolically distinct glucose, ethanol, and quiescent cell states are the starting point in our model construction and a schematic illustration of the dynamic interactions between these states is shown in Fig. 1B.

The dynamics of the different cellular metabolic states cannot be easily observed directly but it is rather straightforward to monitor cell growth by optic growth curve measurements (see Methods). With the help of mathematical modeling, we are able to infer the switching behavior between the metabolic states and the related nutrient dynamics from time-course data. This is done by constructing alternative quantitative growth models with different metabolic switching mechanisms between the states and testing these hypothetical models against time-course data by means of statistical techniques. In the following, we construct a mathematical model that describes yeast cell growth on glucose and ethanol and couples the growth dynamics with transient switching between three distinct metabolic states: (i) glucose, (ii) ethanol, and (iii) quiescent state (Fig. 1B).

We model the cell growth and switching between different metabolic states by ODEs. We start by considering the glucose state in which the cells grow on glucose. We denote the cell mass in this state by *m*^g^. Given that the glucose intake is sufficiently fast, the cell mass dynamics in the glucose state can be modeled as

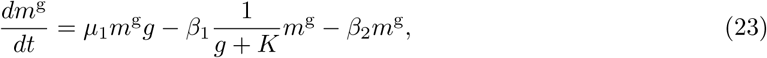

where *g* denotes the level of available glucose and the first term, *µ*_1_*m*^g^*g*, describes the actual growth kinetics with the rate parameter *µ*_1_. If the glucose signal drops to a low level, the cells start to switch gradually to the ethanol state. This switching is modeled using the second term in Eq. 23 and the rate of switching is determined by *β*_1_ and *K*. In a similar manner, the third term in Eq. 23 describes the possible switching to the quiescent state with the rate parameter *β*_2_. In a typical experimental setting, a fixed amount of glucose is provided to cells in the beginning and the glucose level decreases when it is used for growth. Subsequently, the glucose concentration is governed by

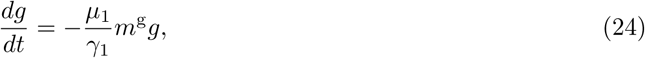

where *γ*_1_ is a parameter that determines the yield of glucose to the produced biomass. Growth in the ethanol state occurs in an analogous manner as in the glucose state. We denote the cell mass in the ethanol state by *m*^e^ and the cell mass dynamics in this state is modeled as

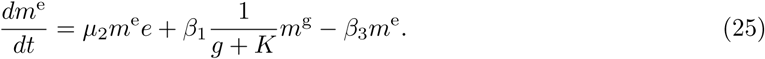

Here, the first term describes the actual growth kinetics with the rate parameter *µ*_2_, the second term corresponds to the cell mass entering the ethanol state from the glucose state, and the third term describes the possible switching from the ethanol state to the quiescent state with the rate parameter *β*_3_. Ethanol is typically not added to a cell culture, but it is produced as a by-product of growth on glucose. Thus, ethanol dynamics is given by

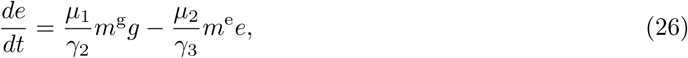

where the first term represents ethanol production during the growth on glucose and the second term considers the decrease due to biomass production. The parameters *γ*_2_ and *γ*_3_ determine the production and decrease, respectively. The above expressions for *m*^g^ and *m*^e^ dynamics include switching to a quiescent state. We denote the cell mass in the quiescent state by *m*^q^ and describe the cell mass dynamics in this state by

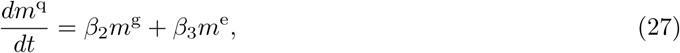

with the terms introduced in Eqs. 23 and 25. Given the three distinct metabolic states, the total cell mass reflecting directly the experimental time-course measurements is given by *m* = *m*^g^ + *m*^e^ + *m*^q^. In experiments, cells are initially put in glucose rich medium and we therefore assume that all cells are initially in the glucose state and the initial glucose level is high. Consequently, we assume that only the model variables *m*^g^ and *g* have non-vanishing initial values. These properties are also used in the reparameterization of the mathematical model which is presented in detail in the Supplementary Information. The model output, i.e. the total cell mass as a function of time, is denoted by *m*(*t, θ*) where *θ* is a parameter vector containing the parameters that result from the reparameterization.

### 3.2 Statistical inference for model parameters and metabolic transitions in homogeneous medium

The mechanisms that are included in the mathematical model are illustrated in Fig. 1B. The full model contains the essential transition from the glucose state to the ethanol state and allows the cells also to switch to the quiescent state directly from the glucose and ethanol states. However, detailed information about the switching mechanisms to the quiescent state is not available and, consequently, there remains notable uncertainty about the routes that cells may use to enter the quiescent state. To treat this uncertainty accurately, we consider three alternative hypotheses (*H*_1_, *H*_2_, and *H*_3_) regarding the switching routes between the metabolic states (schematic illustrations of corresponding switching models are shown in Fig. 1C) and investigate the feasibility of these hypotheses by quantitative statistical testing. In the following, we outline the experimental data used for model calibration and explain how we infer the structure and parameterization of the microenvironment model.

We carried out growth curve measurements for wild-type and petite yeast strains in order to obtain dynamic data on total cell mass that can be used in microenvironment model inference (see Methods). The petite yeast strain differs genetically from the wild-type strain and is not capable to grow on ethanol (DeRisi *et al.*, 1997; Ferea *et al.*, 1999). In the context of our microenvironment model, this means that the growth rate parameter *µ*_2_ should tend to zero when the petite strain is considered but all other parameters can be expected to be shared between these two strains. Given this straightforward connection between the wild-type and petite strains, we can carry out the statistical inference using the wild-type data and subsequently test the predictive performance of our models against the petite strain which is not involved in the model calibration.

We collect the wild-type growth curve data into the data vector *D_k_*. The elements of this data vector contain the average total cell mass at time points *t_k_*, *k* = 1,…, *N*. The average cell mass as well as the corresponding sample variances *v_k_* are computed over 6 replicates (see Supplementary Information for details about data pre-processing). From previous studies (Alvarez-Vasquez *et al.*, 2007; Galdieri *et al.*, 2010; Taylor and Ehrenreich, 2014) the relative fractions of cells in ethanol and quiescent states at steady state (reached in our setting at *t_N_* = 80 hours) can be taken to be approximately 29 ± 6% and 62 ± 6%, respectively. We denote these relative fractions by *α*^e^ = 0.29 and *α*^q^ = 0.62 and the corresponding standard deviations representing uncertainty about the exact values by 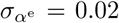 and 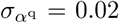. These wild-type data, which are used in model calibration and hypothesis testing, can be combined with the model output under alternative metabolic switching hypothesis *H*_1_, *H*_2_, and *H*_3_ by assuming independent normally distributed measurement errors and defining the likelihood function

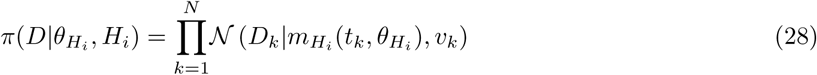

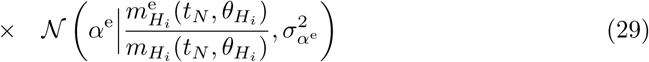

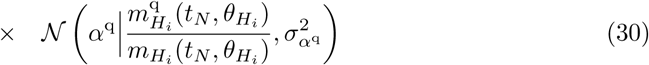

where 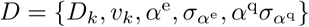 is the data, 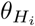 is the parameter vector under the hypothesis *H_i_*, and 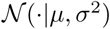 is the normal probability density function with the mean *µ* and variance *σ*^2^. We construct a Bayesian statistical model by combining the likelihood function with uninformative but proper prior distributions where we do not assume any prior dependencies between the parameters and use standard normal prior distributions in logarithmic parameter space. The selected prior distribution introduces a soft lower bound for the parameters. Thus, if a certain rate parameter is present in the model, its value cannot be infinitely close to zero.

### 3.3 Quantitative hypothesis testing reveals the most likely metabolic switching mechanisms

The posterior analysis is first carried out independently for each alternative metabolic switching mechanism (hypotheses *H*_1_, *H*_2_, and *H*_3_). The resulting approximations for the parameter posterior distributions show that the models are identifiable under all three metabolic wiring scenarios (Supplementary Information, Figs. S2-S4, a summary about convergence diagnostics can be found in Figs. S5). In general, the predictions in all three scenarios are in a good agreement with the experimental wild-type data (see predicted total cell mass in Fig. 1C, wild type). The posterior predictive distributions (PPDs) are very similar under the hypotheses *H*_1_ and *H*_2_ and the only notable difference is that under *H*_1_ there is more variability in dynamics (Fig. 1C, wild type). This finding is natural because the models are nested and the additional switching route under hypothesis *H*_1_ increases the model flexibility. The PPD under hypothesis *H*_3_ shows less variability and, further, the dynamic behavior of *m*^e^ is different when compared with the other two scenarios. Further, Fig. 1C shows the PPDs also for the petite strain and we can conclude that under all three hypotheses we are capable of predicting the total cell mass dynamics of the petite strain even though the dynamics of the non-observed model components may differ significantly. Consequently, we can conclude that the predictive performance of our model is good for both the training and the validation data sets. However, based on visual inspection, it is impossible to judge which hypothesis is most likely and, therefore, we carry out statistically rigorous quantitative hypothesis testing over the hypotheses *H*_1_, *H*_2_, and *H*_3_.

Despite the non-distinguishable model predictions in the data space, the posterior analysis over different metabolic switching hypotheses shows significantly more evidence for *H*_2_ (Fig. 1C). The posterior probability of *H*_2_ is very close to one (the posterior probabilities as well as the estimated logarithmic marginal likelihoods are shown in parentheses after the hypothesis name in Fig. 1C). Strong statistical evidence for *H*_2_ suggests that the metabolic switching to the quiescent state in wild-type yeast cells occurs always via the ethanol state in agreement with biological data interpretations (DeRisi *et al.*, 1997; Aragon *et al.*, 2008; Galdieri *et al.*, 2010).

### 3.4 Spatial modeling framework to study colony formation

In our experimental setup, yeast cells grow on a glucose rich agar plate and, in the following, we construct a spatial modeling framework which allows us to predict three dimensional cell state and nutrient distributions during the colony formation process. In addition to cell mass and nutrient dynamics within the colony, we also model the nutrient dynamics within the agar.

To setup the spatial model, we discretize the space into elementary cubes (Fig. 2A). Given that the size of the elementary cubes is chosen appropriately, the growth dynamics within each cube (microenvironment) can be modeled under the homogeneity assumption. In other words, each elementary cube consists of a homogeneous mixture of nutrients and cells in distinct metabolic states (Fig. 2A) and the time-evolution of these local components can be described using the microenvironment model that we constructed above. The spatial colony formation is subsequently determined by the dynamics of interacting neighboring cubes with information coupling between neighboring cubes occurring by the flow of nutrient signals and movement of growing cell mass.

**Figure 2:**
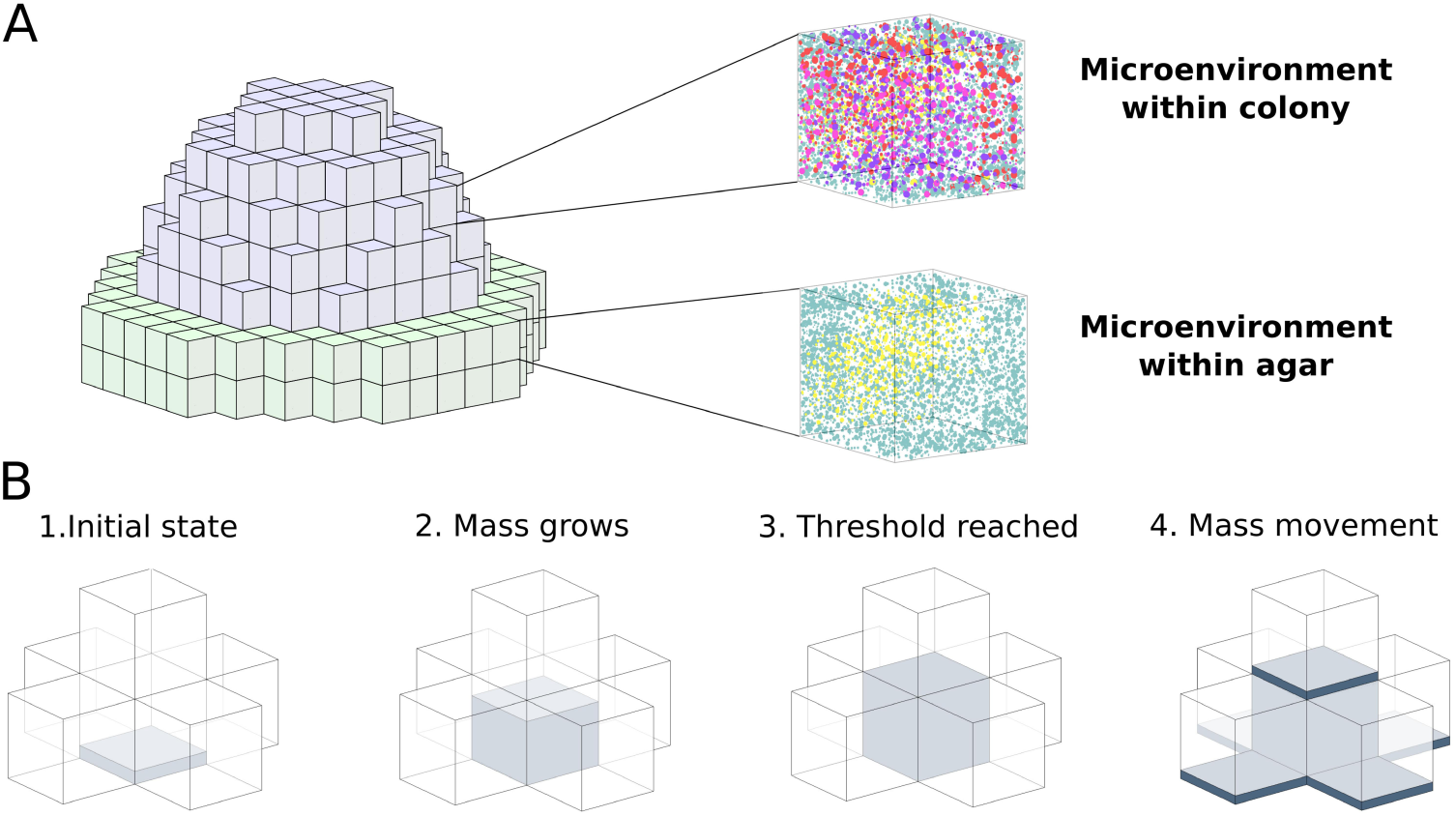
Illustration of the spatial modeling framework. Simulated colonies consist of interacting elementary cubes (for illustrative purposes, the cubes are here notably larger than in practise). (A) Illustration of the elementary cube approximation of a yeast colony. The upper part of the colony (gray elementary cubes) represents the cell mass domain. In these elementary cubes, each microenvironment consists of a mixture of nutrients and cells in different metabolic states. Further, the lower part of the colony (green elementary cubes) represents the nutrient rich agar domain. In the agar domain, each microenvironment can consist of a mixture of nutrients and no cell mass is present. (B) Mass movement is modeled by considering the fill levels of the elementary cubes. The cell mass is growing in the cubes and once a the fill level threshold is reached, cell mass starts to be move into the neighboring cubes. During the cell mass movement, relative fractions of cells in different metabolic states are moved along.

The cell mass movement is modeled by considering fluxes between neighboring cubes. The fluxes are determined by thresholded fill levels of the neighboring cubes and the cell mass is moving from a high concentration to low concentration (for illustration, see Fig. 2B and parameters are given in Tab. 1). The thresholding is essential because the size of elementary cubes is fixed and it is reasonable to assume that the mass movement does not take place until a certain amount of cell mass has accumulated locally and until the pressure starts to push cells forward. In our implementation, the fluxes are computed between six neighboring cubes in each spatial direction and the time-evolution of the full mass distribution is modeled using an ODE system which is determined by the net impact of the individual fluxes. The fluxes are always computed based on the thresholded total mass distribution and the proportions of metabolic states moving along the cell mass are proportional to the proportions of cell states in the cube from which the cell mass is moving. On top of the agar, cell mass can move only to five directions because mass movement into the agar is excluded.

The nutrient transfer is modeled using the same flux-based model as the cell mass movement. However, the thresholding is not needed for the nutrient transfer because it can be assumed that nutrients can diffuse freely over the domain. The domain for glucose diffusion is the union of the agar domain and the elementary cubes with positive cell mass. In addition, it is assumed that the ethanol which is produced as a by-product during growth on glucose can diffuse freely over the positive cell mass. A formal derivation of the mass movement and nutrient transfer models can be found in Methods Section.

### 3.5 Data-driven calibration of the spatial model

As explained in detail above, the spatial model consists of interacting elementary cubes and within each cube we have approximately homogeneous mixture of cells in different metabolic states and nutrients. Local dynamics in each elementary cube are modeled using the microenvironment model whose structure and parameterization is calibrated using growth curve data and population composition information at time 80 hours. More specifically, we use the microenvironment model under metabolic switching hypothesis *H*_2_ which has clearly the highest evidence in statistical testing. The parameterization of the model is fixed to the maximum a posteriori values that were obtained as a by-product of the posterior analysis. Once the microenvironment model is parameterized, we are left with several unknown parameters that are needed for the spatial framework. These parameters are the mass movement rate, the nutrient transfer rates in the agar and within the cell mass, and the initial glucose level in the agar (Table 1). Because practically no pressure is accumulating inside the colony, we set a high value for the mass movement rate (20 h^−1^). This means that the cell mass is distributed at the same rate as the cells are growing and local crowding does not occur. Further, we assume that the glucose reserve in the agar can be modeled by means of a disc with thickness of 0.2 mm and a diameter of 1 cm. Then the local initial glucose level in the elementary cubes in the agar domain can be normalized to equal one and, consequently, we are left with two free parameters: the nutrient transfer rate in the agar and the nutrient transfer rate within the cell mass.

**Table 1:**
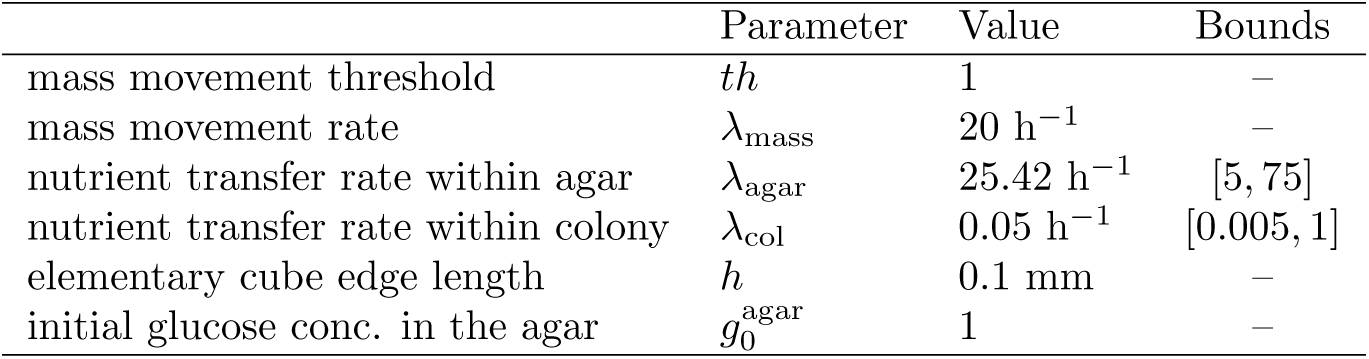
Parameters of the spatial framework. Bounds are given for parameters that are estimated.

To estimate the free parameters of the spatial framework, we measure the area under the growing (wild type) colony over time (for details, see Methods) and optimize the free parameters so that the area under the simulated colony agrees with these data. In other words, we minimize the cost function

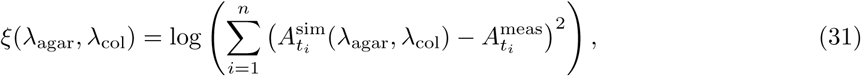

where *λ*_agar_ and *λ*_col_ are the transfer rates within agar and colony, respectively, and 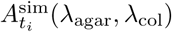 and 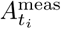 are the simulated and measured areas at time *t_i_*, respectively. Because objective initialization of the cell state and nutrient distribution above the agar is practically impossible, we initialize one elementary cube with cell mass in the glucose state up to the cell mass movement threshold and set the initial glucose level in this cube to one. The simulation is started using this initial state and we assume that a realistic initial state which corresponds the real initial population composition is reached after 5 hours of simulation (this point in time corresponds to the time 0 in the experimental data).

We minimize the cost function using Bayesian optimization (Jones *et al.*, 1998). The optimization is initialized by evaluating the cost function at 20 points which are sampled within the bounds (Table 1) using Latin hypercube sampling. After initialization, the optimal parameter values (Table 1) are obtained after 9 iterations of the algorithm. Fig. 3A shows the fitted footprint area against the experimental data. The model fit is in a good agreement with the data even though at the late time points the model shows saturating behavior that is not present in the real data. This slight disagreement suggests that there is some fraction of cells in a metabolic state which is not included in the model. However, the calibrated model does not only fit well to the wild type data but is also in an excellent agreement with two replicates of our petite strain validation data (see red curves in Fig. 3A). The third replicate can clearly be seen as an outlier and, based on these good fits, we conclude here that our model successfully captures essential dynamics also with respect to the colony size over time.

**Figure 3:**
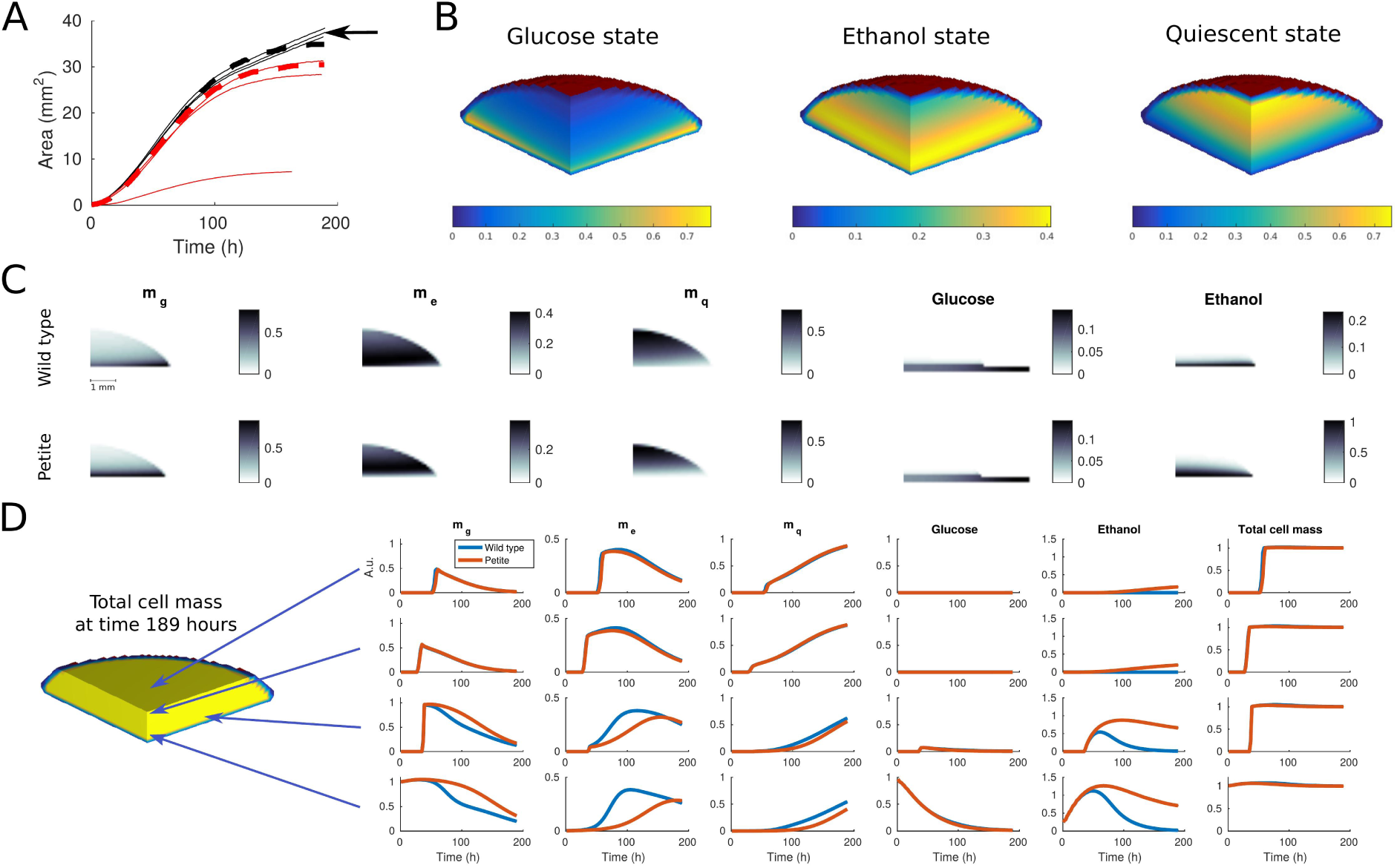
The calibration of the spatial framework and predictions on the colony morphology and colony composition. The colony composition is illustrated for a quarter colony which contains full information of the symmetric colony. (A) Simulated colony footprint areas for wild type and petite strain are plotted using black and red dashed lines, respectively. Experimental data from wild type and petite strains (three replicates from both strains) are plotted using black and red solid lines, respectively. The black arrow indicates the wild type replicate which was used to calibrate the model. The data from the petite strain is used only for validation purposes. (B) Isosurface illustration of the simulated colony shape and cell state composition at time 121 hours. (C) Simulated cell state and nutrient distributions for wild type and petite strains at time 121 hours illustrated using heatmaps. The shown vertical slice is located in the middle of the colony. (D) Simulated time-evolution of all model component all total cell mass at different spatial locations. The exact coornitates (in mm) for illustrated point are (1, 1, 1), (0.1, 0.1, 1.0), (0.1, 1.5, 0.2), and (0.1, 0.1, 0.1) (starting from the upper row).

### 3.6 Predicting nutrient and metabolic state distributions

The calibrated model provides us with rich information about the spatial organization within the colony as well as the colony morphology over time. Fig. 3B illustrates the colony shape and cell state composition at time 121 hours. Interestingly, we see three distinct regions in which different cell states are concentrated. The cells in glucose state are present mainly close to the agar, the cells growing on ethanol are located in the middle of the colony, and, in the upper part the colony, we see a high concentration of cells in quiescent state.

A more detailed view on the spatial organization within the colony is given in Fig. 3C which shows the simulated cell state and nutrient distributions for wild type and petite strains in the middle of the colony at time 121 hours. The nutrient distributions show that glucose is mainly present close to the agar and this indicates that most of the glucose growth and consumption occur in this region. Further, the ethanol distributions show that ethanol level is much higher in the case of the petite strain. This is natural because petite cells produce ethanol but cannot use it for growth and thus there is no consumption. The snapshot distributions for wild type and petite strains look quite similar but essential differences become visible when we observe the time-evolution of model components at different spatial locations (Fig. 3D). Besides the differing ethanol dynamics for wild type and petite strains, also the cell state dynamics differ notably at many spatial locations. The driver for these differences is the growth in the ethanol state which on its behalf affect the switching between the different metabolic states.

## 4 Discussion and conclusions

We introduce a novel coarse-grained multiscale model for yeast colony formation, show how it can be calibrated in a data-driven manner, and validate the calibrated model using independent experimental data. In addition, we illustrate how we are able to predict the spatial organization within the colony as well as the colony morphology over time and discuss the related biological findings.

Even though our computational framework is presented in the context of yeast colony modeling, our approach is fully general and can be applied to model any multicellular system. For instance, an interesting future application for our method could be to study the role of metabolic coupling during human glioblastoma tumor growth. Further, the modeling framework can also be easily extended to study for example the effect of channels that might transport nutrient to different spatial locations within cell mass.

The simulation of our multiscale model is computationally costly and, thus, the calibration of the spatial parameters is a non-trivial task. We formulate the calibration task as an optimization problem in which we minimize the distance between the observed and predicted colony footprint area. In principle, the optimal parameter values could also be found by means of exhaustive parameters sweeps but a notable amount of computational resources can be saved by using Bayesian optimization. The role of this efficient optimization technique becomes even more important when rich data across the scales becomes available and a larger fraction of model parameters can be calibrated together with the spatial parameters.

Our ultimate goal is to develop a spatial framework that would allow simultaneous calibration of local and global parameters. Careful formulation of the related statistical inference problem would also enable at least semi-automatic experimental design planning. In other words, the model calibration could be carried out iteratively so that every iteration would not only provide information about the parameters but also probabilistic predictions on the most beneficial future measurements (e.g. what to measure, when, and at which time point).

## Acknowledgements

The authors acknowledge the computational resources provided by the Aalto Science-IT project.

